# Dickkopf1 is a Novel Endogenous Ligand for priming NLRP3 Inflammasome in Macrophages via TLR4

**DOI:** 10.1101/2025.03.11.642660

**Authors:** Theingi Aung, SuJeong Song, Joyce Kasongo, Hisashi Harada, Shijung Zhang, Octavian Henegariu, Wook-Jin Chae

## Abstract

Dickkopf1(DKK1) is a quintessential Wnt antagonist and immunomodulator in various inflammatory diseases. The underlying molecular mechanisms of DKK1-mediated immunomodulation remain elusive. Here, we identified TLR4 as a new receptor for DKK1 to activate NFκΒ pathway-mediated gene expressions and pyroptosis via NLRP3 inflammasome in human and mouse macrophages.

DKK1 employed TLR4 to initiate NFκB signaling cascade via MyD88. MyD88-TAK1-NFκΒ pathway activation by DKK1 increased HIF1α, NFκB, and NLRP3 protein expression levels, leading to pyroptosis. Unlike LPS, DKK1 did not induce IRAK4 phosphorylation, while the interaction between MyD88 and IRAK4 was maintained for downstream signaling activation. DKK1 did not induce IRF3 phosphorylation in the nucleus and failed to induce IFNβ gene expression, indicating differential signaling from LPS. DKK1 primed macrophages via TLR4-MyD88, resulting in NLRP3 inflammasome-mediated pyroptosis via Caspase-1 and Gasdermin D maturation with various NLRP3 inflammasome activators, including Nigericin. Our results demonstrated that DKK1 is a novel endogenous priming ligand that augments differential NFκB pathway from LPS and NLRP3 inflammasome activation via TLR4 in mouse and human macrophages.

## Introduction

Inflammation plays a crucial role in tissue repair and injury response, acting as both protective and pathological mechanisms. Macrophages are the key immune cells throughout inflammation and repair (Wynn & Barron, 2010). Persistent inflammation caused by dysregulated macrophage activation often leads to chronic inflammation (Valledor et al., 2010).

The Wnt signaling pathway regulates cell differentiation and proliferation for tissue regeneration and wound healing. Dickkopf1 (DKK1), a member of the DKK family proteins, is a quintessential Wnt antagonist known to play a role in inflammation and pathological conditions. Initially identified for its role in embryonic development in *Xenopus laevis*, DKK1 has been implicated in various pathological conditions such as rheumatoid arthritis, pulmonary fibrosis, cardiovascular diseases, Alzheimer’s disease, and cancers (Caricasole et al., 2004; Fezza et al., 2019; Ruaro et al., 2018; Sung, Park et al., 2023a; Toth, 2024).

DKK1’s immunomodulatory role has been characterized in studies using type 2 and type 17 inflammatory disease models (Chae et al., 2016; Wu et al., 2021). By studying DKK1-mediated gene expressions in mouse bone marrow-derived macrophages (mBMDMs) by bulk RNA Seq analysis, it has been reported that DKK1 upregulates proinflammatory genes that are involved in macrophage activation and inflammation (Sung, Song, et al., 2023b). DKK1 binds to Low-density lipoprotein receptor-related protein 6 (LRP6) and inhibits Wnt signaling (Bourhis et al., 2010; S. Chen et al., 2011). With evidence of DKK1’s role in inflammation, a question of the underlying mechanisms of DKK1 to function as an immunomodulator beyond the Wnt signaling inhibitor has been raised, with the possibility of DKK1 binding to its alternative receptor.

The TLR4-MyD88 signaling pathway plays a crucial role in macrophage-mediated innate immune responses. TLR4 requires MD2 and CD14 to form a receptor complex for LPS (lipopolysaccharides) binding (Ciesielska et al., 2021; Nagai et al., 2002). TLR4 recruits MyD88, which interacts with Interleukin-1 receptor-associated kinase 4 (IRAK4). TLR4-MyD88-IRAK4 complex induces the phosphorylation of TGFβ-associated Kinase 1 (TAK1) at Ser 412 upon LPS stimulation, which is crucial for full activation of TAK1 to induce proinflammatory cytokines such as IL-8 (Ouyang et al., 2014). TAK1 phosphorylates IκB Kinase complex (IKKα/β) (De Nardo et al., 2018; Gay et al., 2014; Y. Tan & Kagan, 2019). Activated IKKα/β phosphorylates IκBα which sequesters NFκB p65 in the cytoplasm. Phosphorylated IκBα undergoes degradation and translocates NFκB p65 to the nucleus (Cohen & Strickson, 2017). Activation of NFκB pathway induces NLRP3 and pro-IL-1β gene expressions required for the NLRP3 inflammasome assembly and the subsequent release of IL-1β (Cohen & Strickson, 2017; Gay et al., 2014).

A recent study showed that HIF1α mediates activation of NFκB in hepatic macrophages (X. Wang et al., 2019). HIF1α amplifies inflammation by promoting the proinflammatory signaling pathways. In parallel with canonical NFκB pathway activation, Tank-binding Kinase 1 (TBK1) phosphorylates IRF3 and IRF7 through adaptor molecule TRAM and TRIF by LPS (Fitzgerald et al., 2003; Kagan et al., 2008). The activation of IRF3 and IRF7 is important for Type 1 IFN production, mediating cytotoxic effects and immunostimulatory function in malignant cells or facilitating IL-1β production in endotoxin-challenged mice (Holicek et al., 2024; Sin et al., 2020).

Inflammasomes are cytosolic, multimeric protein complexes mediated by the innate immune system to induce inflammation at the injury site. NLRP3 inflammasome is implicated in various diseases, including atherosclerosis, neurodegenerative diseases, and cancer (Guo et al., 2015; Lamkanfi & Dixit, 2012; Y. Wang et al., 2016; Wei et al., 2014). The activation of NLRP3 inflammasome is a two-step process: priming and activation. Priming (signal 1), triggered by danger signals such as LPS, is recognized by Pattern Recognition Receptor (PRR) such as TLR4, leading to pro-IL-1β synthesis and NLRP3 post-translational modifications. The NFκB pathway acts as a transcriptional activator for NLRP3 and other genes, facilitating inflammasome assembly and activation (Li et al., 2024). Activation (signal 2) is initiated by NLRP3 activators such as Nigericin and MSU, causing inflammasome assembly (Yang et al., 2019). NLRP3 oligomerizes and recruits ASC and pro-caspase-1 upon stimulation, leading to the maturation of Caspase-1. Cleaved caspase-1 processes pro-IL-1β into mature form and also cleaves Gasdermin D. The cleaved Gasdermin D forms membrane pores, thereby causing pyroptosis and secretion of mature IL-1β into extracellular space (W. He et al., 2015; Shi et al., 2015; Swanson et al., 2019). In this study, we investigated the mechanism of DKK1-mediated signaling for proinflammatory gene expressions via the TLR4-MyD88-TAK1-NFκB pathway in macrophages. We demonstrated that the DKK1-mediated NFκB signaling pathway primes macrophages to induce NLRP3 inflammasome activation, which plays a crucial role in multiple inflammatory diseases.

## Results

### DKK1 induces proinflammatory gene expressions via the NFκB pathway in macrophages

Recent studies indicated that DKK1 acts as an immunomodulator (Chae & Bothwell, 2019). Our previous work indicated that DKK1 induces proinflammatory gene expressions in murine macrophages such as *Il1b*, *Hif1α*, *Nlrp3*, and *Relb*, involving NFκB pathway activation (Sung, Song, et al., 2023b). However, how the proinflammatory signaling cascade was activated by DKK1 in macrophages remains elusive. To investigate the molecular mechanisms of DKK1-mediated gene expressions, we decided to use the human monocytic THP1 cell line.

We used LPS as a positive control in our experiments and compared it with DKK1, given that LPS induces inflammation by upregulating IL-1β, NLRP3, and HIF1α protein expressions in macrophages (Q. Chen et al., 2022; Gicquel et al., 2015). THP1 cells were stimulated with PMA for 48 hours for differentiation, followed by LPS or DKK1 treatment for 4 and 24 hrs. DKK1 and LPS induced NLRP3 and pro-IL-1β expressions at 4 hr, and their upregulations sustained till 24 hr. HIF1α expression was upregulated by DKK1 treatment at 24 hr, similar to LPS, but with little expression at 4 hr, suggesting that DKK1 upregulates proinflammatory gene expressions in macrophages (**Fig 1A**).

**Figure 1.**
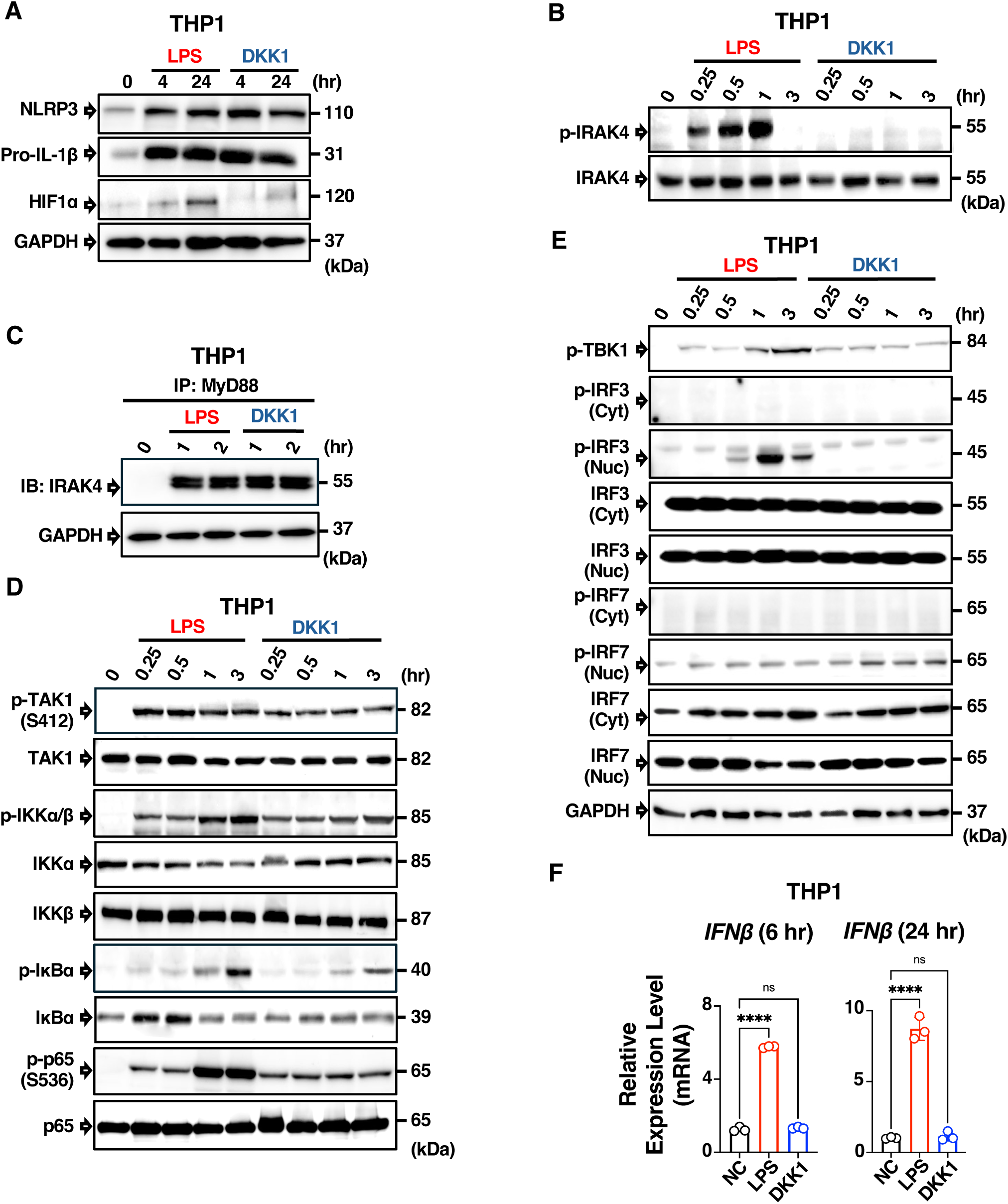
DKK1 induces pro-inflammatory gene expressions via NFκB pathway in macrophages. **(A)** Western blot analysis of WT THP1 macrophages after treated with Vehicle control (NC), 100 ng/ml LPS or 30 ng/ml DKK1 for 4 hr and 24 hr and probed with anti-HIF1α, anti-NLRP3, anti-pro-IL-1β antibodies. GAPDH was used as control. **(B)** Western blot analysis of WT THP1 macrophages treated with Vehicle control (NC), 100 ng/ml LPS or 30 ng/ml DKK1 for 15 min, 30 min, 1 hr, 3 hr and probed with anti-p-IRAK4, anti-IRAK4 antibodies **(C)** WT THP1 macrophages were treated with Vehicle control (NC), 100 ng/ml LPS or 30 ng/ml DKK1 for 1 hr and 2 hr. Cell lysates were immunoprecipitated with anti-MyD88 Ab and immunoblotted with anti-IRAK4 Ab. GAPDH was used as a loading control. **(D)** Western blot analysis of WT THP1 macrophages treated with Vehicle control (NC), 100 ng/ml LPS or 30 ng/ml DKK1 for 15 min, 30 min, 1 hr, 3 hr and probed with anti-p-TAK1^Ser412^, anti-TAK1, anti-p-IKKα/β, anti-IKKα, anti-IKKβ, anti-p-IκBα, anti-IκBα, anti-p-p65^Ser536^, anti-p65 antibodies. **(E)** Western blot analysis of WT THP1 macrophages treated with Vehicle control (NC), 100 ng/ml LPS or 30 ng/ml DKK1 for 15 min, 30 min, 1 hr, 3 hr and whole cell lysates were probed with anti-TBK1 and anti-GAPDH antibodies. Cytoplasmic and Nuclear extracts were probed with anti-p-IRF3, anti-IRF3, anti-p-IRF7 and anti-IRF7 antibodies. **(F)** THP1 macrophages were treated with Vehicle control (NC), 100 ng/ml LPS or 30 ng/ml DKK1 for 6 hr or 24 hr. Relative expression levels of human *IFNβ* mRNA were normalized to *GAPDH* (when *GAPDH* is 10). Statistically significant differences were analyzed by one-way ANOVA and Bonferroni’s multiple comparisons test. (ns, not significant, **** p<0.0001) A representative of two independent experiments is shown.

Recognition of pathogen-associated molecular patterns (PAMPs) initiates the recruitment of MyD88 and IRAK4 (Kawai & Akira, 2010). IRAK4 is the upstream kinase in the NFκB signaling pathway and regulates the stability of the myddosome complex with MyD88 to induce proinflammatory responses in mouse and human macrophages (Ferrao et al., 2014; Kawagoe et al., 2008; Kawai & Akira, 2010). IRAK4 undergoes autophosphorylation, which is required for its kinase activity (Cheng et al., 2007). Unlike LPS, DKK1 did not induce IRAK4 phosphorylation, and we confirmed that IRAK4 phosphorylation did not occur by DKK1 up to 24 hr (**Fig 1B, Fig EV1A**). Given that IRAK4 scaffold with MyD88 mediates proinflammatory cytokine production via NFκB and MAPK regardless of IRAK4 kinase activity in human macrophages (Cushing et al., 2017; Pereira et al., 2022; Qin et al., 2004; J. Sun et al., 2016), we tested the interaction between IRAK4 and MyD88 without IRAK4 phosphorylation. Immunoprecipitation (IP) by MyD88 Ab indicated that IRAK4 interacts with MyD88 upon DKK1 stimulation (**Fig 1C**). Reverse IP analysis using IRAK4 Ab confirmed MyD88-IRAK4 interaction (**Fig EV1B**).

Next, we examined TAK1^Ser412^ phosphorylation, which is crucial for proinflammatory cytokine production in response to LPS via NFκB (Ouyang et al., 2014). TAK1 is a key mediator of MyD88-driven signaling and strongly correlated to the activation of NFκΒ (Qian et al., 2001). DKK1 induced phosphorylation of TAK1^Ser412^ and its downstream kinase, IKKα/β (**Fig 1D**). IκBα was phosphorylated upon DKK1 treatment, similar to LPS. Consistent with these results, p65^Ser536^ phosphorylation was observed (**Fig 1D**).

LPS stimulation phosphorylates TBK1, activating IRF3 and IRF7 to induce transcription of type 1 IFN genes (Sakaguchi et al., 2003; Sin et al., 2020; Y. Tan & Kagan, 2019). DKK1 phosphorylated TBK1, but the phosphorylation levels were weaker than those induced by LPS. Unlike LPS, DKK1 did not induce IRF3 phosphorylation, while it did phosphorylate IRF7 in the nucleus (**Fig 1E**). Consistent with Fig 1E, IFNβ mRNA expressions were induced by LPS but not by DKK1, indicating that DKK1-mediated TBK1 and IRF7 activation was insufficient to induce IFNβ gene expressions (**Fig 1F**). Taken together, our data suggested that DKK1 utilizes differential NFκB pathway signaling from LPS.

### DKK1 utilizes MyD88 and TAK1 to activate NFκB pathway

We reasoned that DKK1-induced NFκB activation utilizes MyD88 as an adaptor protein. To test this hypothesis, we utilized MyD88 KO THP1 macrophages. MyD88 deficiency in THP1 macrophages was confirmed by Western blot (**Fig EV2A**). NLRP3 protein expressions were diminished in MyD88 KO THP1 macrophages upon DKK1 treatment. Little pro-IL-1β and HIF1α protein expressions were detected in MyD88 KO THP1 macrophages upon DKK1 stimulation, suggesting MyD88’s important role in inducing these protein expressions (**Fig 2A**). Consistently, TAK1^Ser412^, IKKα/β, IκBα, and p65^Ser536^ phosphorylation were markedly diminished by MyD88 deficiency (**Fig 2B**). MyD88 ablation markedly abolished TBK1 phosphorylation in the cytosol and reduced IRF7 phosphorylation in the nucleus (**Fig 2B, Fig EV2B**). These data indicated that DKK1 utilizes MyD88 as an adaptor protein to activate NFκΒ signaling pathways.

**Figure 2.**
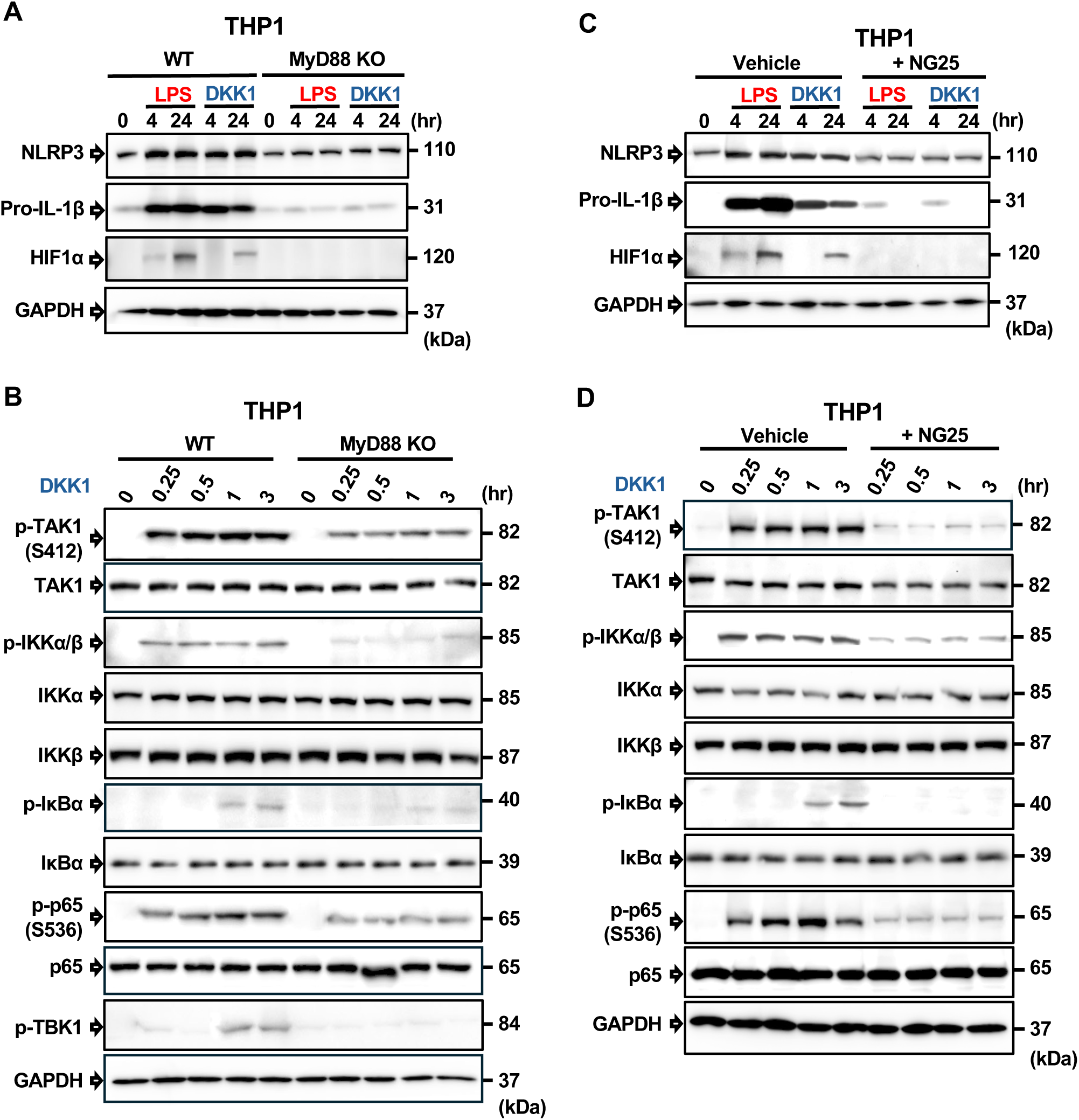
DKK1 utilizes MyD88 and TAK1 to activate NFκB pathway. **(A)** Western blot analyses of WT and *MyD88* KO THP1 macrophages after treated with Vehicle control (NC), 100 ng/ml LPS or 30 ng/ml DKK1 for 4 hr and 24 hr and probed with anti-HIF1α, anti-NLRP3, anti-pro-IL-1β antibodies. GAPDH was used as loading control. **(B)** Western blot analysis of WT and *MyD88* KO THP1 macrophages treated with Vehicle control (NC) or 30 ng/ml DKK1 for 15 min, 30 min, 1 hr, 3 hr and probed with anti-p-TAK1^Ser412^, anti-TAK1, anti-p-IKKα/β, anti-IKKα, anti-IKKβ, anti-p-IκBα, anti-IκBα, anti-p-p65 ^Ser536^, anti-p65, anti-p-TBK1, anti-GAPDH antibodies. **(C)** WT THP1 macrophages were pretreated with 10 μM NG25 for 1 hr, followed by 100 ng/ml LPS or 30 ng/ml DKK1 for 4 hr, 24 hr and probed with anti-HIF1α, anti-NLRP3, anti-pro-IL-1β, anti-GAPDH antibodies. **(D)** WT THP1 macrophages were treated with 30 ng/ml DKK1 for 15 min, 30 min, 1 hr, 3 hr after pretreated with or without 10 μM NG25 for 1 hr. Cell lysates were probed with anti-p-TAK1 ^Ser412^, anti-TAK1, anti-p-IKKα/β, anti-IKKα, anti-IKKβ, anti-p-IκBα, anti-IκBα, anti-p-p65^Ser536^, anti-p65, anti-GAPDH antibodies. A representative of two independent experiments is shown.

We decided to test the role of TAK1 in the DKK1-induced NFκB signaling pathway. HIF1α, NLRP3, and pro-IL-1β protein expressions were probed by pre-treatment of a TAK1 inhibitor (NG25) and DKK1 treatment (L. Tan et al., 2015). NLRP3 protein expressions were reduced, and HIF1α protein expression was undetectable by TAK1 inhibition. Pro-IL-1β protein expression gradually decreased by 24 hr by TAK1 inhibition (**Fig 2C**). We confirmed a significant inhibition of TAK1^Ser412^ phosphorylation by TAK1 inhibitor (**Fig 2D**). Further analyses on DKK1-mediated NFκB pathway activation revealed that TAK1 inhibition subsequently decreased the phosphorylation of IKKα/β, IκBα, and p65^Ser536^, suggesting an important role of TAK1 in DKK1-mediated NFκB pathway activation (**Fig 2D**).

### DKK1 requires TLR4 as a receptor for the NFκB signaling pathway activation

The interaction between DKK1 and LRP6 was characterized (S. Chen et al., 2011). We have observed that ablation of LRP6 in myeloid lineage cells reduced inflammation (Sung et al., 2024). We anticipated that the ablation of LRP6 in myeloid lineage cells reduces DKK1-mediated gene expressions in mBMDMs. qPCR analyses of these genes upon DKK1 stimulation in mBMDMs were performed using *Lrp6* ^fl/fl^ and *LysMCre*-*Lrp6* ^fl/fl^ mice. Unexpectedly, we found that LRP6 ablation did not show a marked decrease of *Hif1α*, *Arg1*, *Cd274*, and *Marco* gene expressions, while *Il1r1* and *Il1β* mRNA only partially decreased upon DKK1 treatment, suggesting that LRP6 is not the primary receptor of DKK1-induced proinflammatory gene expressions (**Fig 3A**).

**Figure 3.**
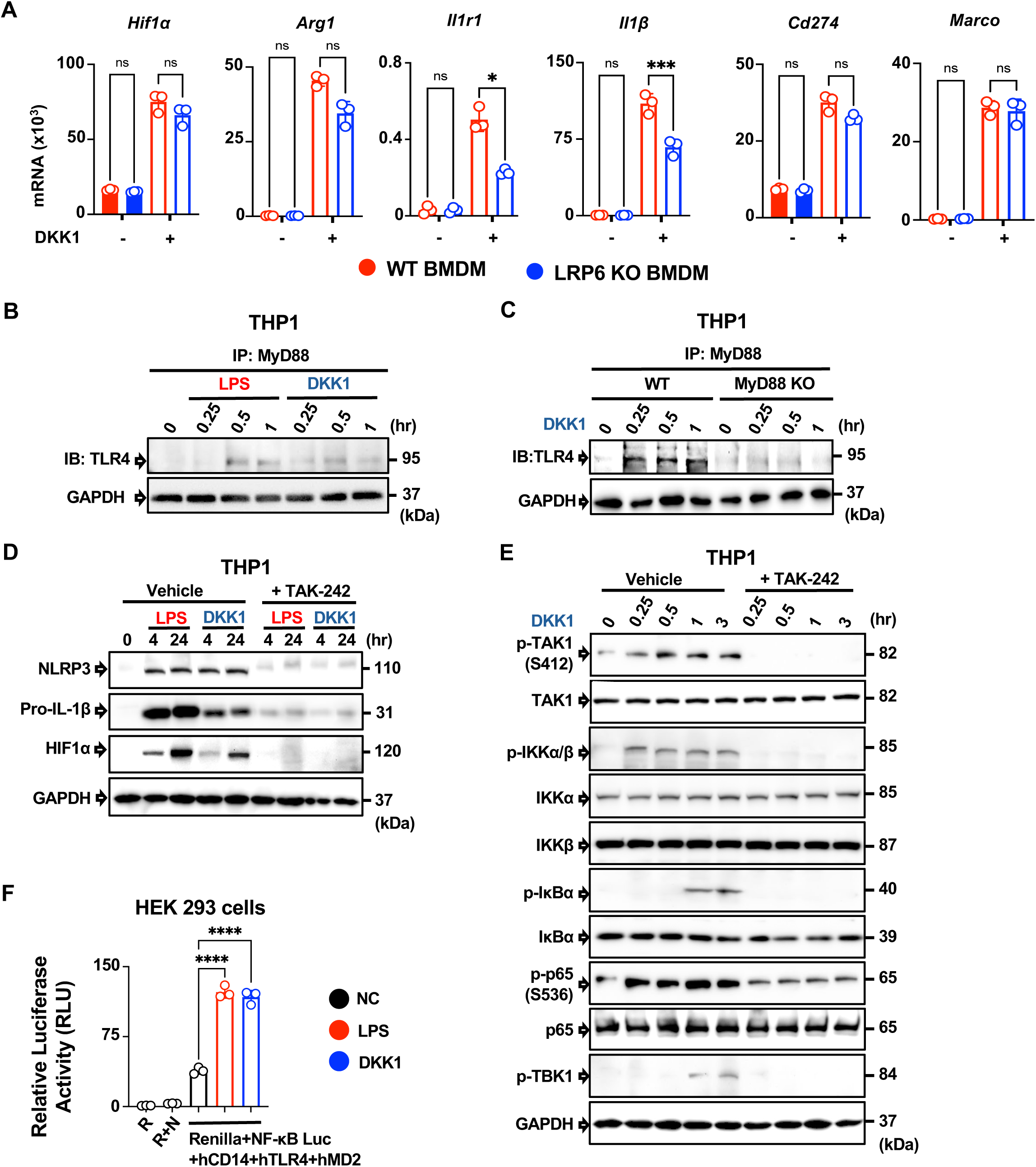
DKK1 requires TLR4 as a receptor. **(A)** mBMDM from WT and and *LysMCre-Lrp6 ^fl/fl^* mice were treated with Vehicle control (NC) or 30 ng/ml DKK1 for 6 hr and *Hif1α, Arg1, Il1r1, Il1β, Cd274 and Marco* mRNA levels were measured by qPCR. *Tbp* was used as housekeeping gene. **(B)** WT THP1 macrophages were treated with Vehicle control (NC), 100 ng/ml LPS or 30 ng/ml DKK1 for 15 min, 30 min and 1 hr. Cell lysates were immunoprecipitated with anti-MyD88 antibody and immunoblotted with anti-TLR4 antibody. GAPDH was used as a loading control. **(C)** WT and *MyD88* KO THP1 macrophages were treated with Vehicle control (NC) or 30 ng/ml DKK1 for 15 min, 30 min and 1 hr. Cell lysates were immunoprecipitated with anti-MyD88 antibody and immunoblotted with anti-TLR4 antibody. GAPDH was used as a loading control. **(D)** WT THP1 macrophages were pretreated with 100 nM TAK-242 for 1 hr, followed by 100 ng/ml LPS or 30 ng/ml DKK1 for 4 hr, 24 hr and probed with anti-HIF1α, anti-NLRP3, anti-pro-IL-1β and anti-GAPDH antibodies. **(E)** WT THP1 macrophages were treated with 30 ng/ml DKK1 for 15 min, 30 min, 1 hr, 3 hr after pretreated with or without 100 nM TAK-242 for 1 hr. Cell lysates were probed with anti-p-TAK1^Ser412^, anti-TAK1, anti-p-IKKα/β, anti-IKKα, anti-IKKβ, anti-p-IκBα, anti-IκBα, anti-p-p65^Ser536^, anti-p65, anti-p-TBK1, anti-GAPDH antibodies. **(F)** HEK 293 cells were transfected with plasmids encoding hTLR4, hMD2 and hCD14 cDNAs. NFkB luciferase activity was measured after stimulating with 20 ng/ml LPS or 75 ng/ml DKK1 for 10 hr using *Renilla* luciferase (pRL-CMV) as a control. Statistically significant differences were analyzed by one-way ANOVA and Bonferroni’s multiple comparisons test (ns, not significant, * p<0.05, *** p<0.001). A representative of two independent experiments is shown.

Our results led us to hypothesize that DKK1 may need a receptor that requires MyD88 to activate the NFκB signaling pathway. We hypothesized that TLR4 may be an alternative receptor of DKK1. To answer this question, we tested the DKK1-induced interaction of TLR4 and MyD88 by IP in THP1 macrophages. Flow cytometry and western blot analyses confirmed that THP1 macrophages stably expressed TLR4, CD14, and MD2 proteins (**Fig EV3A, EV3B**).

Immunoprecipitation using MyD88 Ab and immunoblot with TLR4 Ab was performed upon DKK1 stimulation at each time point. The interaction between TLR4 and MyD88 was detected upon DKK1 stimulation by 30 min, similar to LPS (**Fig 3B**). We confirmed that TLR4-MyD88 interaction did not occur in MyD88 KO THP1 macrophages upon DKK1 treatment like LPS, validating the interaction between TLR4 and MyD88 upon DKK1 treatment (**Fig 3C, Fig EV3C**).

To further investigate DKK1-TLR4 downstream signaling, we used a well-known TLR4 inhibitor, TAK-242, and analyzed DKK1-induced protein expressions and NFκB pathway activation (Matsunaga et al., 2011). NLRP3 and pro-IL-1β protein expressions were reduced, and HIF1α protein expression was undetectable by TAK-242 treatment (**Fig 3D**). Next, we investigated DKK1-mediated NFκB pathway inhibition by TAK-242. TAK1^Ser412^ phosphorylation clearly disappeared by TAK-242. Consistently, the phosphorylation of IKKα/β, IκBα, and p65^Ser536^ were decreased by TAK-242, indicating that DKK1 requires TLR4 to induce NFκB pathway activation (**Fig 3E**). In addition, DKK1-mediated TBK1 phosphorylation was notably diminished by TLR4 inhibition (**Fig 3E**).

As TLR4 functions together with MD2 and CD14 in the recognition of LPS (Akashi et al., 2003), we measured NFκB-mediated gene transcription activity in HEK 293 cells transfected with plasmids encoding hTLR4, hMD2, and hCD14 cDNAs. Stimulation of transfected HEK 293 cells with LPS or DKK1 induced a similar level of NFκB luciferase activity, suggesting that TLR4 works together with its co-receptors MD2 and CD14 upon DKK1 stimulation (**Fig 3F**). Collectively, we demonstrated that TLR4 is a receptor of DKK1 to induce the NFκB pathway activation.

### DKK1 acts as a primer for pyroptosis

We showed that DKK1-mediated NFκB pathway activation induces the upregulation of NLRP3 and IL-1β protein expressions. We hypothesized that DKK1 acts as a primer for pyroptosis and IL-1β secretion. First, we measured the percentage of cell death by Lactate Dehydrogenase (LDH) release and secretion of IL-1β in mBMDMs and THP1 macrophages. mBMDMs and THP1 macrophages were stimulated with LPS or DKK1 as primers at each dose, followed by Nigericin as a second signal to activate NLRP3 inflammasome for DKK1-induced pyroptosis and IL-1β secretion in mBMDMs and THP1 macrophages (**Fig 4A, 4B**). Next, we tested DKK1-mediated pyroptosis with different activation signals, such as SiO2 nanoparticles (Nano-SiO_2_) and Monosodium Urate crystal (MSU). Like LPS, DKK1 induced pyroptosis and ΙL-1β secretion in mBMDMs and THP1 macrophages (**Fig 4C, 4D**). These data suggested that DKK1 primes macrophages for pyroptosis and IL-1β secretion.

**Figure 4.**
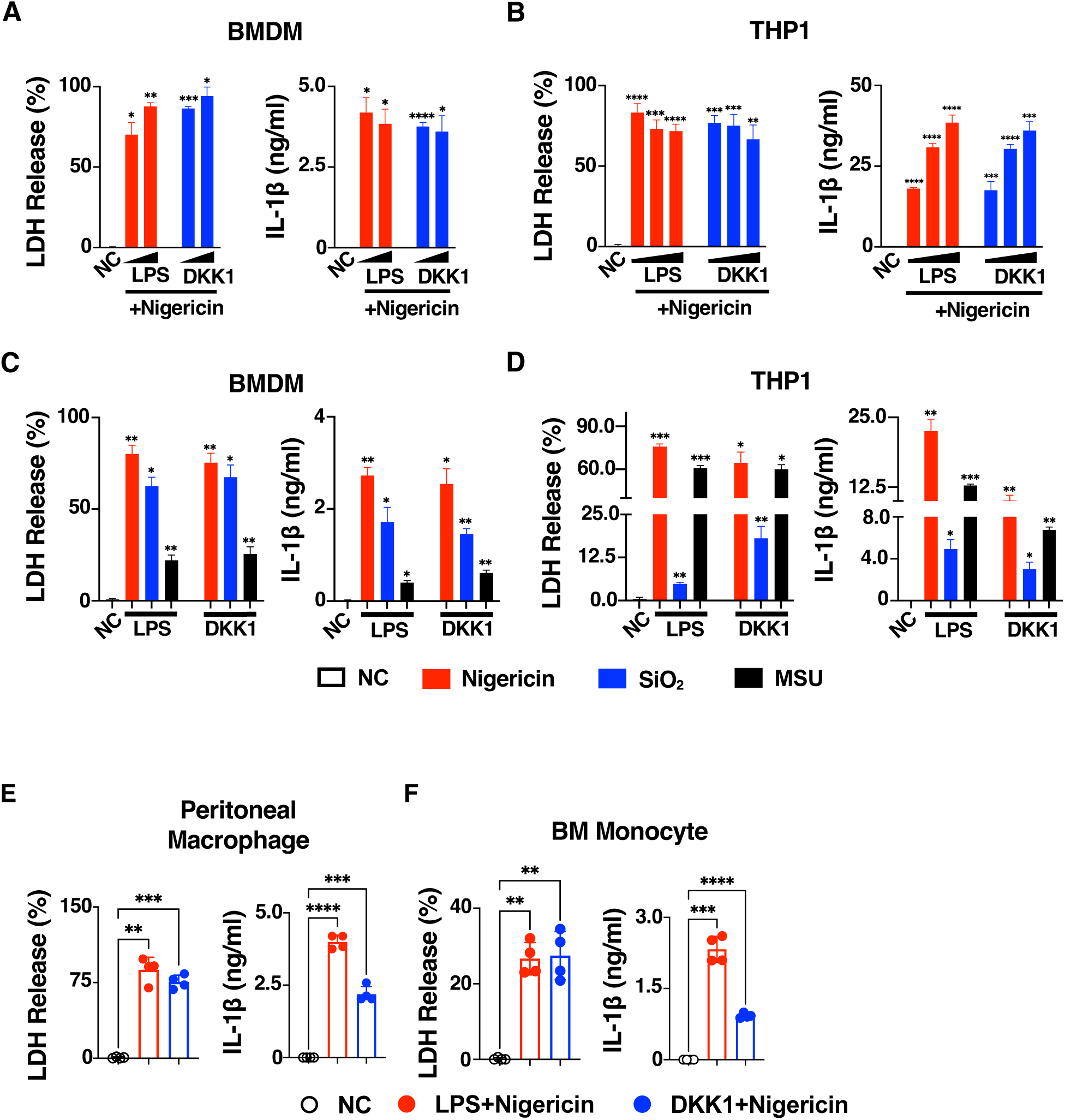
DKK1 acts as a primer for pyroptosis. **(A)** Mouse BMDM were treated with Vehicle control (NC), LPS (100, 1000 ng/ml) or mouse DKK1 (30, 100 ng/ml) for 3 hr followed by 10 μM Nigericin for 1.5 hr. **(B)** THP1 macrophages were treated with Vehicle control (NC), LPS (1, 10, 100 ng/ml) or human DKK1 (3, 30, 100 ng/ml) for 3 hr followed by 10 μM Nigericin for 1.5 hr. **(C, D)** Mouse BMDMs and THP1 macrophages were treated with Vehicle control (NC), 100 ng/ml LPS or 30 ng/ml DKK1 for 3 hr, followed by 10 μM Nigericin for 1.5 hr, 250 μg/ml Nano-SiO_2_ for 6 hr or 300 μg/ml MSU for 6 hr. (**E, F, G**) Peritoneal Macrophages, bone marrow monocytes from mice, and *in vitro*-generated MDSCs were treated with Vehicle control (NC), 100 ng/ml LPS or 30 ng/ml DKK1 for 3 hr, followed by 10 μM Nigericin for 1.5 hr. Pyroptotic cell death was measured by LDH release and IL-1β secretion was measured by ELISA. Statistically significant differences were analyzed by one-way ANOVA and Bonferroni’s multiple comparisons test. (* p<0.05, ** p<0.005, *** p<0.001, **** p<0.0001). A representative of two independent experiments is shown.

DKK1 induced pyroptosis and IL-1β secretion in peritoneal macrophages and mouse bone marrow-derived monocytes, indicating that DKK1 induces pyroptosis in other types of macrophages (**Fig 4E, 4F**). Collectively, our data suggested that DKK1 acts as a primer to induce pyroptosis and IL-1β secretion in mouse and human macrophages.

### DKK1 primes NLRP3 inflammasome-induced pyroptosis

NLRP3 inflammasome activation, a crucial component of the innate immune system, is linked to pyroptosis (W. He et al., 2015). As DKK1 acted as a primer to induce pyroptosis and IL-1β secretion, we investigated whether DKK1 priming leads to NLRP3 inflammasome activation-mediated pyroptosis. We used MCC950, an inhibitor that directly binds to NLRP3 domains and suppresses its activation (Coll et al., 2015). MCC950 markedly decreased the levels of pyroptosis and IL-1β release in mBMDMs and THP1 macrophages (**Fig 5A, 5B**).

**Figure 5.**
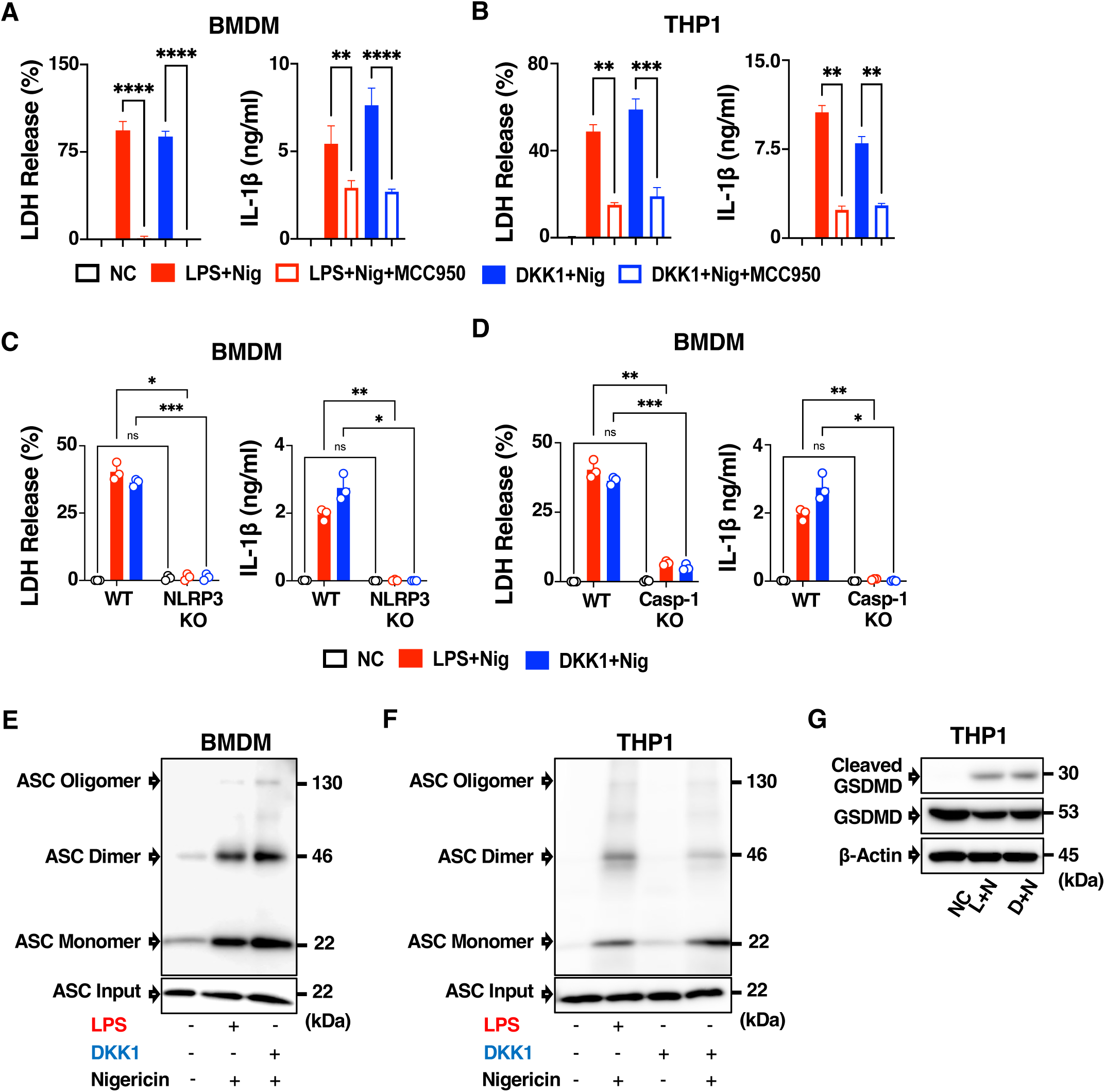
DKK1 primes NLRP3 inflammasome-induced pyroptosis. **(A, B)** LDH release and IL-1β secretion were investigated in cell supernatants of mouse BMDMs **(A)** and THP1 macrophages **(B)** after treatment with Vehicle control (NC), LPS/DKK1 and Nigericin with or without 0.5 μM MCC950. **(C)** LDH release and IL-1β secretion were investigated in WT vs *Nlrp3* KO BMDM after treatment with Vehicle control (NC), LPS or DKK1 with Nigericin **(D)** LDH release and IL-1β secretion were investigated in WT vs *Caspase-1* KO BMDM after treatment with Vehicle control (NC), LPS or DKK1 with Nigericin. **(E, F)** Western blot analysis of ASC oligomerization in mouse BMDMs **(E)** and THP1 macrophages **(F)** after stimulating with Vehicle control (NC), 100 ng/ml LPS or 30 ng/ml DKK1 for 3 hr, followed by 10 μM Nigericin for 1.5 hr. **(G)** Western blot analysis of Gasdermin D and Cleaved Gasdermin D in THP1 macrophages after stimulating with Vehicle control (NC), 100 ng/ml LPS or 30 ng/ml DKK1 for 3 hr, followed by 10 μM Nigericin for 1.5 hr. β-Actin was used as a loading control. Statistically significant differences were analyzed by one-way or two-way ANOVA and Bonferroni’s multiple comparisons test (ns, not significant, * p<0.05, ** p<0.005, *** p<0.001, **** p<0.0001). A representative of two independent experiments is shown.

To test whether DKK1 utilizes NLRP3 and Caspase-1 to induce pyroptosis, we measured pyroptosis and IL-1β secretion in *Nlrp3*- and *Caspase-1* KO mice-derived BMDMs. *Nlrp3*- and *Caspase-1* deficiency substantially decreased pyroptosis and IL-1β release, indicating that DKK1 utilizes these two conventional NLRP3 inflammasome pathway components (**Fig 5C, 5D, Fig EV4A, 4B**). Consistently, cleavage of pro-IL-1β was abolished in *Nlrp3-* and *Caspase-1*-KO mBMDMs, indicating the important role of NLRP3 and Caspase-1 in DKK1-mediated IL-1β maturation (**Fig EV4A, 4B**).

The oligomerization of ASC (Apoptosis speck-like protein containing CARD) and the formation of ASC-SPECK are one of the key indicators of NLRP3 inflammasome assembly (Dick et al., 2016). We tested DKK1-mediated ASC oligomerization in mBMDMs and THP1 macrophages. ASC oligomerization did occur upon LPS or DKK1, followed by Nigericin treatment in mBMDMs and THP1 macrophages (**Fig 5E, 5F**). Immunofluorescence microscopy further confirmed ASC-SPECK formation (**Fig EV4C**). Cleavage of Gasdermin D determines pyroptosis by releasing cleaved Gasdermin D-N-terminal (GSDMD-NT) with intrinsic pyroptosis-inducing activity (Shi et al., 2015). Gasdermin D was cleaved by DKK1 priming followed by Nigericin treatment (**Fig 5G**). Taken together, our data suggested that DKK1 induces pyroptosis and IL-1β secretion via canonical NLRP3 inflammasome activation.

### DKK1 utilizes TLR4 and MyD88 to augment pyroptosis and IL-1β secretion

We showed that the TLR4-MyD88-TAK1-NFκB pathway is important to induce DKK1-mediated NLRP3 protein expression. LRP6 deficiency reduced inflammation in myeloid lineage cells (Sung et al., 2024). We first tested whether DKK1 requires LRP6 for pyroptosis. Consistent with qPCR data from *LysMCre-Lrp6* ^fl/fl^ mice in Fig 3A, LRP6 ablation showed little effect on pyroptosis and IL-1β secretion by DKK1, confirming that LRP6 is dispensable (**Fig 6A**). We used a DKK1 inhibitor (WAY-262611), which inhibits DKK1’s effect on the canonical Wnt signaling pathway (Chae et al., 2016; Pelletier et al., 2009; Wu et al., 2021). DKK1 inhibitor did not decrease pyroptosis in THP1 macrophages, suggesting that DKK1 does not utilize the canonical Wnt signaling pathway for pyroptosis (**Fig 6B**).

**Figure 6.**
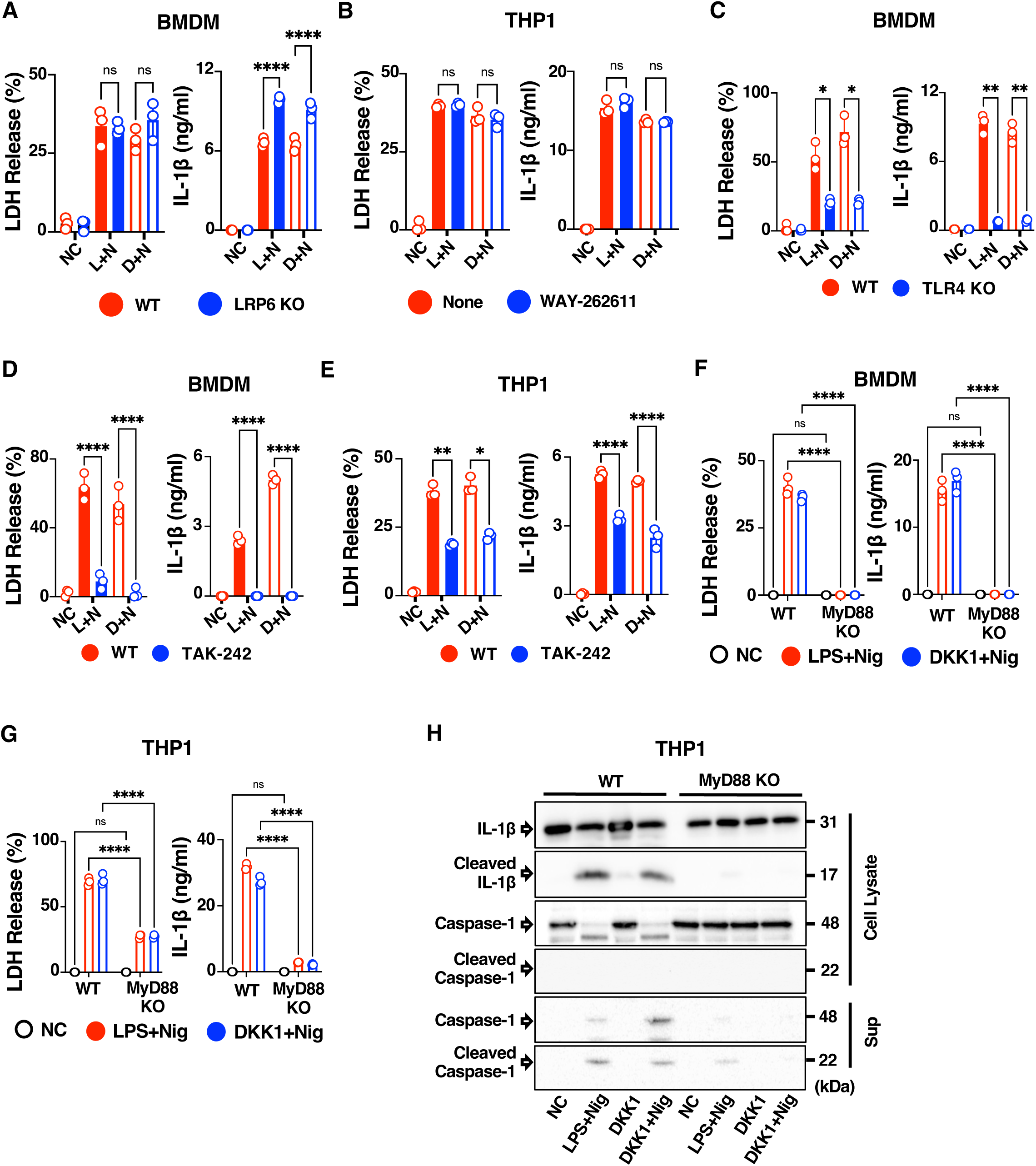
DKK1 utilizes TLR4 and MyD88 to augment NLRP3 inflammasome-induced pyroptosis. **(A)** BMDMs from WT and *LysMCre-Lrp6 ^fl/fl^* mice were treated with 100 ng/ml LPS or 30 ng/ml DKK1 for 3 hr, followed by 10 μM Nigericin for 1.5 hr. **(B)** WT THP1 macrophages were pretreated with 2 μM WAY-262611 for 1 hr, followed by treatment as in **(A)**. **(C)** BMDMs from WT and *Tlr4* KO mice were treated as in (**A**). **(D, E)** BMDMs from WT mice and WT THP1 macrophages were treated as in **(A)** with or without 100 nM TAK-242 for 1 hr. **(F, G)** WT and *MyD88* KO mBMDMs **(F)** and THP1 macrophages **(G)** were treated as in **(A)**. Cell death and IL-1β secretion were measured by LDH assay and ELISA, respectively. **(H)** Western blot analysis of IL-1β, cleaved IL-1β, caspase-1 and cleaved caspase-1 in WT vs *MyD88* KO THP1 cell lysates or supernatants after stimulating with Vehicle control (NC), 100 ng/ml LPS or 30 ng/ml DKK1 for 3 hr, followed by 10 μM Nigericin for 1.5 hr. Statistically significant differences were analyzed by one-way or two-way ANOVA and Bonferroni’s multiple comparisons test (ns, not significant, * p<0.05, ** p<0.005, *** p<0.001, **** p<0.0001). A representative of two independent experiments is shown.

As DKK1 employed TLR4-MyD88 for NLRP3 protein expression, we used *Tlr4* KO mBMDMs to measure DKK1-mediated pyroptosis and IL-1β secretion. TLR4 deficiency markedly decreased pyroptosis and IL-1β secretion by DKK1 or LPS, followed by Nigericin treatment (**Fig 6C**). We further confirmed that TAK-242-mediated inhibition of TLR4 signaling decreased pyroptosis and IL-1β secretion in mBMDMs and THP1 macrophages (**Fig 6D, 6E**). These results indicated that DKK1 utilizes TLR4 as a receptor, activating the NFκB pathway to induce pyroptosis and IL-1β secretion.

To investigate MyD88’s role in DKK1-mediated NLRP3 inflammasome activation, we used MyD88-deficient THP1 macrophages and mBMDMs. MyD88 ablation significantly inhibited pyroptosis and IL-1β release in LPS-or DKK1-primed cells following Nigericin treatment (**Fig 6F, 6G**). Maturations of pro-IL-1β and pro-caspase-1 were blocked by MyD88 deficiency (**Fig 6H, Fig EV5A**). We confirmed that DKK1-induced Gasdermin D cleavage requires MyD88 (**Fig EV5B**). These results indicated that MyD88 is required for NLRP3 inflammasome response upon DKK1 priming.

We used pharmacological inhibitors to examine DKK1-induced NFκB and TAK1 activation for NLRP3 inflammasome activation. mBMDMs and THP1 macrophages were pre-treated with BAY 11-7802, SN50 and NG25 to inhibit NFκB and TAK1. Pyroptosis and IL-1β secretion were inhibited in mBMDMs and THP1 macrophages (**Fig EV5C, 5D**).

A previous study demonstrated an important role of TBK1 in promoting LPS-primed NLRP3 inflammasome activation (Cui et al., 2024). To test the role of TBK1 in DKK1-mediated pyroptosis, we used GSK8612, which binds to TBK1 and inhibits the downstream effects of TBK1 activation (Thomson et al., 2019). DKK1-induced pyroptosis and IL-1β secretion were only partially decreased by TBK1 inhibition (**Fig EV5E**). We further explored if other pathways are involved. We tested the mTOR (mammalian Target Of Rapamycin) pathway, PI3K/Akt (Phosphoinositide 3-kinase/protein kinase B) pathway, SGK1 (Serum and Glucocorticoid-Regulated Kinase 1) pathway, and the Wnt pathway in mBMDMs using Rapamycin, LY294002, GSK653094 and TWS119, respectively. Pyroptosis and IL-1β secretion were not decreased, suggesting that the DKK1-induced NLRP3 inflammasome activation pathway requires a TAK1-NFκB pathway (**Fig EV5F**). Collectively, our results demonstrated that DKK1 utilizes the TLR4-MyD88-TAK1-NFκB pathway for NLRP3 inflammasome-mediated pyroptosis and IL-1β secretion (**Fig 7**).

**Figure 7.**
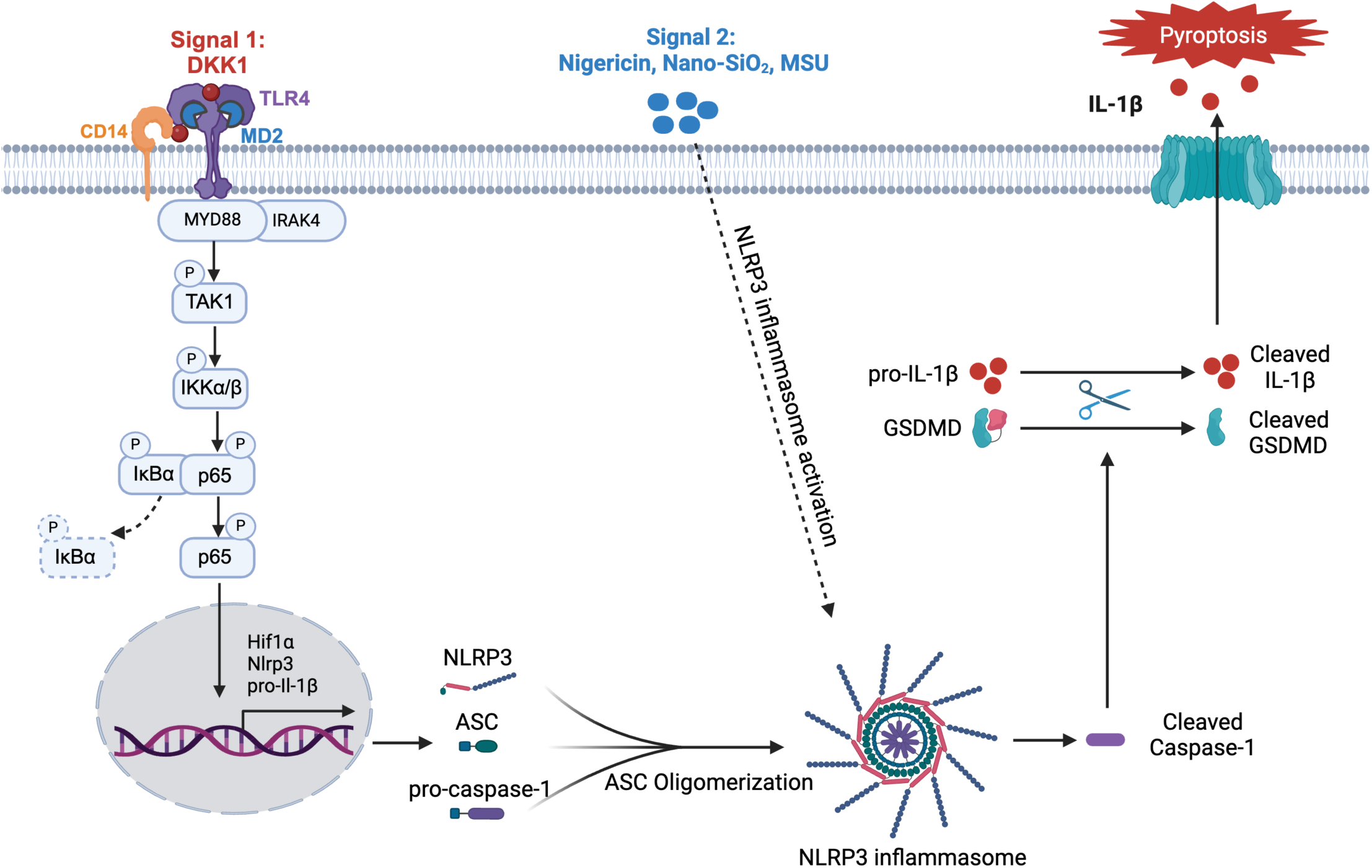
DKK1-mediated NFκB signaling pathway augments NLRP3 inflammasome-induced pyroptosis and IL-1β secretion via TLR4. DKK1 binds to TLR4, recruits MyD88 and IRAK4 to induce TAK1^Ser412^, IKKα/β, IκBα and p65^Ser536^ phosphorylation. DKK1-mediated NFκB pathway activation primes macrophages to upregulate NLRP3, pro-IL-1β and HIF1α protein expressions. Activation signals (Nigericin, Nano-SiO_2_, MSU) induces NLRP3 inflammasome activation, leading to ASC oligomerization and recruiting caspase-1. NLRP3 inflammasome activation cleaves pro-Caspase-1, which further cleaves pro-IL-1β and GSDMD. Cleaved GSDMD forms membrane pores and releases IL-1β via pyroptosis.

## Discussion

In this study, we showed that DKK1 activates NFκB signaling pathways via TLR4-MyD88-TAK1, acting as an endogenous primer for NLRP3 inflammasome activation in mouse and human macrophages.

LRP6 has been studied as a putative receptor for the Wnt signaling inhibition by DKK1.The immunological roles of DKK1 in multiple preclinical models raised the question of the alternate receptor for DKK1 in immune cells. The limited use of LRP6 by DKK1 for pyroptosis provides a novel insight that DKK1 modulates innate immune responses via a core immune sensor such as TLR4 to propagate its downstream signaling pathways rather than using the canonical Wnt signaling pathway.

Our findings indicated that a differential targeting strategy needs to be considered from blocking LRP6 *per se* to control DKK1-mediated pyroptosis and IL-1β secretion. For instance, we and others previously showed that DKK1 inhibitor (WAY-262611) attenuates HDM-induced asthma and *Candida albicans* infection in the lung, yet pyroptosis and IL-1β secretion were not inhibited by WAY-262611 in the current study (Chae et al., 2016; Wu et al., 2021).Since tissue remodeling and repair processes require LRP6 via the canonical Wnt pathway activation, our study will provide a distinctive way to target the DKK1-TLR4 axis in inflammatory diseases. DKK1 has been implicated in various human diseases and murine preclinical models with its elevated levels in local inflamed tissues or circulating levels. Further studies on the role of the DKK1-TLR4 axis in DKK1-high chronic inflammatory diseases are warranted.

Previously, other TLR4 or MD2-binding ligands were reported, such as HMGB1 and Paclitaxel, with different binding modes from LPS requiring a co-receptor complex with CD14 and MD2 (M. He et al., 2018; Paudel et al., 2019; Resman et al., 2014; S. Sun et al., 2019; Szajnik et al., 2009). We showed that DKK1 induces NFκB pathway via TLR4, MD2, and CD14, while the outcome of downstream signaling is different from LPS. DKK1-mediated NFκB pathway is more specialized for the canonical NFκB pathway activation rather than type I IFN induction.

The lack of phosphorylation in IRAK4 and IRF3, as well as the lack of IFNβ expression, upon DKK1 stimulation implicates that DKK1 is differentially recognized from LPS by TLR4 in the host. For example, the lack of IRAK4 phosphorylation suggested that additional host proteins may be recruited to MyD88 upon DKK1 stimulation, promoting effector responses. A previous study by Sun et al. demonstrated that TLR4 signaling in human macrophages relies on IRAK1 activity for IL-6 and TNFα production rather than IRAK4 (J. Sun et al., 2016). DKK1 did not induce IL-6 and TNFα mRNA expressions in our previous study (Sung, Song, et al., 2023b). Further studies are warranted on the detailed DKK1-TLR4 interaction mechanism using the structural biology approach and its downstream signaling.

We and others previously demonstrated that platelets are circulating sources of DKK1 (Chae et al., 2016; Kagey & He, 2017; Sung, Song, et al., 2023b; Ueland et al., 2009). In addition, cancer cells also express DKK1 (Malladi et al., 2016), suggesting that the dysregulated DKK1 expression levels or participation of activated platelets in the affected organs can be attributed to the increased levels of DKK1 in pathological inflammation. Our study highlighted DKK1 as a primer for NLRP3 inflammasome activation in combination with various activation signals, implicating a significant link between DKK1 and NLRP3 inflammasome-mediated pyroptosis to promote chronic inflammatory diseases.

Using TLR4 by DKK1 might shed light on evolutionarily conserved mechanisms of the Wnt signaling and innate immune signaling across species. Hemocytes from *Drosophila Melanogaster* harbors Toll receptors and function for hemostasis, while DKK1 was initially found *in Xenopus laevis*, implying that the Wnt system and innate immune system were diverged in mammals to mount more specific and coordinated immune responses. DKK1 is also released upon infection pathogens such as influenza virus (IAV) or *Candida albicans* infections, indicating that the duration, intensity, and cellular niche of DKK1 expression determines the disease outcome. Future studies on increased DKK1 expression levels, sources of DKK1, and the role of the DKK1-TLR4 axis using disease models and human specimens are required to understand DKK1-mediated inflammatory disorders.

In conclusion, our findings demonstrate that DKK1 triggers the TLR4-MyD88-TAK1-NFκB pathway and acts as a novel endogenous priming ligand for the NLRP3 inflammasome-mediated pyroptosis, placing the DKK1-TLR4 axis as an important target for regulating inflammation.

## Materials and Methods

### Mice

C57Bl/6J mice (#000664), MyD88 KO mice (#009088), Caspase-1 KO mice (#016621), Nlrp3 KO mice (#021302), Tlr4 KO mice (#029015) were purchased from the Jackson Laboratory and have been bred in our mouse facility. *LysMCre*-*Lrp6* ^fl/fl^ mice were generated as previously described (Sung et al., 2024). All mouse protocols were approved by the VCU Animal Care and Use Committee (IACUC) approval in accordance with the Association for Assessment and Accreditation of Laboratory Animal Care International (AAALAC).

### Generation of Bone Marrow-derived Macrophages (BMDM)

Generation of BMDM from mice was established in our previous paper (Sung et al., 2024; Sung, Park, Song, et al., 2023). Briefly, femurs and tibias from 6-8 weeks-old mice were isolated, and bone marrow was resuspended in Red Blood Cell lysis buffer (Biolegend, Cat# 420302) to lyse RBCs. After centrifugation at 1400 rpm for 5 min at 4°C, bone marrow cells were resuspended in PBS. The cells were passed through a 40 μm strainer to obtain a single-cell suspension. Cells were resuspended in DMEM supplemented with 20% FBS, 1% Pen/Strep, 1X Glutamax, 1% MEM-NEAA, 1% Sodium Pyruvate and 20 ng/ml M-CSF. For differentiation, cells were seeded at 4×106/well and cultured for 6 days at 37°C. mBMDMs were used on Day 7 with RPMI-1640 media supplemented with 10% FBS, 1% Pen/Strep, 1X Glutamax, 1% MEM-NEAA, 1% sodium pyruvate and 5 ng/ml MCSF.

### Isolation of Bone Marrow-derived Monocytes and peritoneal macrophages

Bone marrow cells were prepared as above and resuspended in MojoSort buffer. Bone marrow-derived monocytes were isolated using Mouse BM-derived monocytes Mojosort kit (Biolegend, Cat#480153) based on the manufacturer’s protocol. Peritoneal lavage was collected from the peritoneal cavity by injecting PBS. The collected peritoneal lavage was centrifuged at 1400 rpm at 4°C for 5 min. The cell pellet was resuspended in DMEM supplemented with 10% FBS, 1% Pen/Strep, 1X Glutamax, 1% MEM-NEAA, and 1% Sodium Pyruvate.

### Cell Culture

Wild-type THP1 monocytic cells (Invivogen, Cat#t hp-null) and MyD88 KO THP1 cells (Invivogen, Cat# thpd-komyd) were maintained in RPMI-1640 media supplemented with 10% FBS, 1% Pen/Strep, 1X Glutamax, 0.05 mM 2-mercaptoethanol at 37°C with 5% CO_2_. ΗΕΚ 293 cells were cultured in DMEM medium supplemented with 10% FBS, 1% Pen/Strep, and 1X Glutamax.

### LDH Assay

For macrophage differentiation of THP1 cells, cells were treated with 100 nM phorbol 12-myristate-13-acetate (SelleckChem, Cat# 7791) for 24 hr. For priming, THP1 cells were treated with 100 ng/ml LPS and 30 ng/ml hDKK1 for 3 hr, followed by 10 μM of Nigericin for 1.5 hr. For mBMDM, mDKK1 was used. For untreated cells (NC), complete media with DMSO and ethanol was used as a vehicle control. For positive controls, cells were exposed to 1X lysis buffer (ThermoFisher Scientific, Cat# 20300) for 45 min. Cell death in mBMDMs and THP1 macrophages was measured using CyQUANT^TM^ LDH Cytotoxicity Assay kit (Invitrogen, Cat# C20300) according to the manufacturer’s instructions. Inhibitor information is available in the Reagents and Tools table.

### ELISA

IL-1β secreted from mBMDMs and THP1 macrophages were evaluated using ELISA MAXTM Deluxe Set Mouse IL-1β (Biolegend, Cat# 432604) and ELISA MAXTM Deluxe Set Human IL-1β (Biolegend, Cat# 437004) according to the manufacturer’s instructions.

### Luciferase assay

HEK 293 cells were plated and co-transfected with NFκB luciferase reporter using lipofectamine 3000 (ThemoFisher, Cat# L3000001) along with hTLR4, hMD2, hCD14, and *Renilla* luciferase constructs. Cells were treated with LPS and DKK1 after 42 hr of transfection. Plasmid information is listed in the Reagents and Tools table. Luciferase assays were performed 10 hr after LPS/DKK1 treatment under the Dual-Glo® Luciferase assay System (Promega, Cat# E2920), using Renilla Luciferase (pRL-CMV) as a control.

### Western blot

THP1 cells were seeded at 2×10^6^ cells/well and differentiated with 10 nM PMA for 48 hr. THP1 and mBMDMs cell lysates were collected using 1X Lysis buffer prepared from 10X CST Cell Lysis Buffer, Protease inhibitor cocktail, Phosphatase inhibitor cocktail, and Phenylmethylsulfonyl Fluoride (PMSF). Nuclear and cytoplasmic extracts were collected using NE-PER Nuclear and Cytoplasmic Extraction Reagents (ThermoFisher, Cat# 78833) supplemented with 1X Protease Inhibitors, 1X Phosphatase Inhibitors cocktails, and 1X PMSF. Bicinchoninic acid (BCA) assay was performed for protein quantification using PierceTM BCA Protein Assay Kit (Thermo Fisher Scientific, Cat# 23225). For immunoblots, the PVDF membranes were immunoblotted with primary antibodies at 1:2000 dilution at 4°C overnight, after blocking with 5% nonfat milk/TBST for 1 hr. After incubation with anti-rabbit HRP-conjugated secondary antibody in 1:4000 dilution for 1 hr at RT, the membrane was exposed to 1X SignalFire ECL reagent, and images were detected by Chemidoc^TM^ Imaging System (Bio-Rad). Antibodies used for immunoblots are listed in the Reagents and Tools table.

### ASC Oligomerization Assay

Wild type and MyD88 KO THP1 cells were differentiated with 100 nM PMA for 24 hr. The cells were stimulated with LPS (100 ng/ml) and DKK1 (30 ng/ml) for 3 hr, followed by Nigericin (10 μM) for 1 hr. Cells were lysed with 1X RIPA buffer containing 250 mM Tris-HCL (pH 7.4), 750 mM NaCl, 5% NP-40, 2.5% Sodium Deoxycholate, 0.5% SDS supplemented with protease inhibitor cocktail for 20 min on ice and centrifuged at 12000 rpm for 20 min at 4 °C. The cell pellets were washed with PBS and cross-linked with 2 mM Disuccinimidyl Suberate (DSS) in PBS for 30 min at RT in rotation at 225 rpm. After centrifugation at 300 g for 10 min at 4 °C, the cell pellets were resuspended in 1X SDS buffer and boiled at 95 °C for 5 min. ASC oligomerization was visualized using an anti-ASC/TMS1 antibody.

### Immunofluorescence

Wild type and MyD88 KO mBMDMs were seeded at 1×104 cells/well in the 8-well chamber (ThermoFisher, Cat# 154941pk) and stimulated with LPS (100 ng/ml) and mDKK1 (30 ng/ml) for 3 hr followed by Nigericin (10 μM) for 1 hr. The cells were fixed in 4% Paraformaldehyde (PFA) for 15 min and permeabilized with 0.3% Triton X-100 in PBS for 10 min. After blocking in 3% BSA/PBS for 30 min in a humidified chamber, the cells were incubated with anti-ASC/TMS1 Ab in 1:800 dilution in the humidified chamber at 4°C overnight. The next day, the cells were incubated with Alexa Fluor 568-conjugated goat anti-rabbit IgG for 1 hr at RT. The cells were washed, mounted (VECTASHIELD Vibrance Antifade Mounting Medium with DAPI), air dried, and visualized by Olympus Inverted Fluorescence Microscope (Olympus, IX71).

### Immunoprecipitation

WT and MyD88 KO THP1 cells were seeded at 2×106 cells/well and differentiated with 10 nM PMA for 48 hr. Cell lysates were collected using 1X Lysis buffer prepared from 10X CST Cell Lysis Buffer, Protease inhibitor cocktail, Phosphatase inhibitor cocktail, and PMSF. Cell lysates were precleared with 10 μl of protein A magnetic beads for 30 min at RT with rotation for immunoprecipitation. 500 μg of protein was immunoprecipitated with anti-MyD88 Ab in 1:100 dilution and incubated with rotation at 4°C overnight. The next day, 20 μl of Protein A Magnetic Beads were added to each sample. After incubating for 20 min at RT with rotation, the immunocomplex was washed five times with 500 μl of 1X CST cell lysis buffer. Magnetic beads with immunocomplex were mixed with 1X SDS buffer and boiled at 95°C for 5 min. Western blot was proceeded by probing anti-TLR4 polyclonal Ab, anti-MyD88 Ab, and anti-IRAK4 Ab.

### Quantitative real-time PCR

Total RNA was extracted from mBMDMs and THP1 macrophages using TRIzol reagent according to the manufacturer’s instructions, followed by cDNA preparation using a High-Capacity RNA-to-CDNA Kit (Thermofisher Scientific, Cat#4387406). Quantitative real-time PCR was performed using PowerUp SYBR Green master mix with the Applied Biosystems StepOnePlus Real-Time PCR system (Thermofisher Scientific). Primer pairs for human and mouse genes are listed in the Reagents and Tools table.

### Flow Cytometry

THP1 cells were plated in 1×10^6^ cells and differentiated with 100 nM PMA for 24 hr. After blocking a human FC Receptor Blocking Ab (ThermoFisher, Cat# 14-9161-73), the cells were stained with Zombie Aqua Fixable Viability Kit and then incubated with fluorescent-conjugated antibodies against TLR4 and CD14. The data were acquired using FACSymphonyA1 and analyzed by FlowJo software (version 10.7, Tree Star).

### Statistical Analysis

Data are presented as mean values, and SD is indicated by error bars if not indicated. GraphPad Prism software (version 9.4.1, GraphPad Software Inc.) was used for statistical analysis. Statistical significance was determined by one-way or two-way ANOVA analysis with Bonferroni’s multiple comparisons test. P values of <0.05 were considered statistically significant. Data are representative of at least two independent experiments.

## Acknowledgment

We thank Dr. Paula Bos and Dr. Shijung Zhang (Virginia Commonwealth University) for providing insights and critical reading of the manuscript.

## CRediT authorship contribution statement

**Theingi Aung:** Data curation; formal analysis; investigation; methodology; validation; visualization; writing-original draft; writing-review and editing; conceptualization**. Sujeong Song**: Data curation; formal analysis; investigation; methodology; validation; visualization; writing-review and editing. **Joyce Kasongo**: Data curation; methodology; validation; visualization; writing-review and editing. **Hisashi Harada:** Writing-review and editing; validation investigation; methodology. **Shijun Zhang**: Writing-review and editing; validation investigation; methodology. **Octavian Henegariu:** Validation investigation; methodology. **Wook-Jin Chae:** Data curation; formal analysis. funding acquisition; investigation; methodology; project administration; resources; software; supervision; validation; visualization; writing-original draft; writing-review and editing; conceptualization.

