## Supplementary information for "Dickkopf1 is a Novel Endogenous Ligand for priming NLRP3 Inflammasome in Macrophages via TLR4"

**Expanded View Figure Legends**

**Figure EV1. DKK1 maintains MyD88-IRAK4 interaction without IRAK4 phosphorylation (A)** WT THP1 macrophages were treated with medium (NC) or 30 ng/ml DKK1 for 15 min, 30 min, 1 hr, 2 hr, 4 hr, 24 hr and probed with anti-p-IRAK4, anti-IRAK4 and anti-GAPDH antibodies. **(B)** WT THP1 macrophages were immunoprecipitated with anti-IRAK4 antibody and probed with anti-MyD88 antibody. GAPDH was used as a loading control. A representative of two independent experiments is shown.

**Fig EV2. DKK1-induced IRF7 phosphorylation requires MyD88 (A)** Western blot analysis of WT and *MyD88* KO THP1 macrophages probed with anti-MyD88 antibody. **(B)** Western blot analysis of WT and *MyD88* KO THP1 macrophages treated with Vehicle control (NC) or 30 ng/ml DKK1 for 15 min, 30 min, 1 hr and 3 hr. Cytoplasmic and nuclear extracts were analyzed by western blot and probed with anti-p-IRF7 and anti-IRF7 antibodies. A representative of two independent experiments is shown.

**Fig EV3. TLR4 interacts with MyD88 in response to LPS (A)** Flow cytometry analysis of cell surface expression of TLR4 and CD14 in THP1 macrophages using Isotype Ab (blue) and anti-TLR4 Ab, anti-CD14 Ab (red). **(B)** Western blot analysis of WT THP1 macrophages probed with anti-MD2 antibody. **(C)** WT and *MyD88* KO THP1 macrophages were treated with Vehicle control (NC) or 100 ng/ml LPS for 15 min, 30 min and 1 hr. Cell lysates were immunoprecipitated with anti-MyD88 antibody and immunoblotted with anti-TLR4 antibody. GAPDH was used as a loading control. A representative of two independent experiments is shown.

**Fig EV4. DKK1 utilizes NLRP3, Caspase-1 and ASC to induce NLRP3 inflammasome activation (A)** WT vs *Nlrp3* KO mBMDMs were treated with Vehicle control (NC), 100 ng/ml LPS or 30 ng/ml DKK1 for 3 hr, followed by 10 µM Nigericin for 1.5 hr. Whole cell lysates were probed with anti-NLRP3 and anti-cleaved IL-1β antibodies. **(B)** WT vs *Caspase-1* KO mBMDMs were treated with Vehicle control (NC), 100 ng/ml LPS or 30 ng/ml DKK1 for 3 hr, followed by 10 µM Nigericin for 1.5 hr. Whole cell lysates were probed with anti-Caspase-1 and anti-cleaved IL-1β antibodies. **(C)** Immunofluorescence images of ASC-SPECK formation in mBMDMs after treated with Vehicle control (NC), 100 ng/ml LPS or 30 ng/ml DKK1 for 3 hr, followed by 10 μM Nigericin for 1.5 hr. ASC-SPECK (red) are indicated by arrow. Scale bar indicates 50 μM. A representative of two independent experiments is shown.

**Fig EV5. DKK1 utilizes MyD88-TAK1-NFκB pathway to induce pyroptosis and IL-1β secretion (A)** Western blot analysis of IL-1β, cleaved IL-1β, caspase-1 and cleaved caspase-1 in WT vs *MyD88* KO mBMDMs cell lysates after stimulating with Vehicle control or 30 ng/ml DKK1 for 3 hr, followed by 10 μM Nigericin for 1.5 hr**. (B)** Western blot analysis of Gasdermin D and Cleaved Gasdermin D in WT and *MyD88* KO THP1 macrophages after stimulating with Vehicle control (NC), 100 ng/ml LPS or 30 ng/ml DKK1 for 3 hr, followed by 10 μM Nigericin for 1.5 hr. β-Actin was used as a loading control. **(C, D)** mBMDMs and THP1 macrophages were pretreated with BAY 11-7802, NG25 and SN50 for 1 hr. The cells were treated with 100 ng/ml LPS or 30 ng/ml DKK1 for 3 hr, followed by 10 μM Nigericin for 1.5 hr. **(E)** WT THP1 macrophages were pretreated with 20 μM GSK8612 for 1 hr. The cells were treated with 100 ng/ml LPS or 30 ng/ml DKK1 for 3 hr, followed by 10 μM Nigericin for 1.5 hr. **(F)** mBMDMs were pre-treated with different inhibitors for 1 hr followed by DKK1+Nigericin treatment. For inhibitors, Rapamycin, LY 294002, GSK 653094, TWS119 were used. Cell death and IL-1β secretion were measured by LDH assay and ELISA, respectively. Statistically significant differences were analyzed by one-way ANOVA and Bonferroni's multiple comparisons test (ns, not significant, * $p<0.05$, ** $p<0.005$, *** $p<0.001$, **** $p<0.0001$). A representative of two independent experiments is shown.

#### Reagents and Tools Table

| Reagent/Resource | Reference or Source | Identifier or Catalog Number |
|---|---|---|
| **Experimental Models** | | |
| **Mouse models** | | |
| C57Bl/6J (*M. musculus*) | Jackson Lab | 000664 |
| B6.129P2(SJL)-*Myd88*$^{tm1.1Defr}$/J | Jackson Lab | 009088 |
| B6N.129S2-*Casp*$^{tm1Flv}$/J | Jackson Lab | 016621 |
| B6N.129S6-*Nlrp3*$^{tm1Bhk}$/J | Jackson Lab | 021302 |
| B6(Cg)-*Tlr4*$^{tm1.2Karp}$/J | Jackson Lab | 029015 |
| *LyzM*Cre-*LRP6*$^{fl/fl}$ | (Sung et al., 2024) | |
| **Cell lines** | | |
| THP1 | Invivogen | thp-null |
| *MyD88* KO THP1 | Invivogen | thpd-komyd |
| HEK 293 cells | Dr. Iain Morgan (VCU School of Dentistry) | |
| **Recombinant DNA** | | |
| Plasmid pRL-CMV (Renilla Luciferase) | Promega | E2261 |
| Plasmid NFκB-Luc | Agilent | 219077 |
| pcDNA3-human CD14 | Addgene | 13645 |
| human-TLR4 cDNA wt pDEST40 | Addgene | 42646 |
| pFlag-CMV1-human MD2 | Addgene | 13028 |
| **Antibodies** | | |

| | | |
|---|---|---|
| Anti-HIF1α | Cell Signaling Technology | 36169 |
| Anti-NLRP3 | Cell Signaling Technology | 15101 |
| Anti-pro-IL-1β (mouse) | Cell Signaling Technology | 31202 |
| Anti-pro-IL-1β (human) | Cell Signaling Technology | 12703 |
| Anti-Cleaved IL-1β (mouse) | Cell Signaling Technology | 63124 |
| Anti-Cleaved IL-1β (human) | Cell Signaling Technology | 83186 |
| Anti-pro-Caspase-1 | Cell Signaling Technology | 83383 |
| Anti-Cleaved Caspase-1 | Cell Signaling Technology | 9661 |
| Anti-Gasdermin D | Cell Signaling Technology | 39754 |
| Anti-Cleaved Gasdermin D | Cell Signaling Technology | 36425 |
| Anti-p-p65$^{Ser\ 536}$ | Cell Signaling Technology | 3033 |
| Anti-NFκB p65 | Cell Signaling Technology | 8242 |
| Anti-p-IKKα/β | Cell Signaling Technology | 2697 |
| Anti-IKKα | Cell Signaling Technology | 61294 |
| Anti-IKKβ | Cell Signaling Technology | 8943 |

| | | |
|---|---|---|
| Anti-p-IκBα | Cell Signaling Technology | 2859 |
| Anti-IκBα | Cell Signaling Technology | 4812 |
| Anti-p-TAK1$^{Ser\ 412}$ | Cell Signaling Technology | 9339 |
| Anti-TAK1 | Cell Signaling Technology | 5206 |
| Anti-p-TBK1 | Cell Signaling Technology | 5483 |
| Anti-GAPDH | Cell Signaling Technology | 5174 |
| Anti-TLR4 | ThermoFisher | 48-2300 |
| Anti-β-Actin | Cell Signaling Technology | 4970 |
| Anti-ASC/TMS1 | Cell Signaling Technology | 13833 |
| Anti-MyD88 | Cell Signaling Technology | 4283 |
| Anti-p-IRF3 | Cell Signaling Technology | 29047 |
| Anti-IRF3 | Cell Signaling Technology | 11904 |
| Anti-p-IRF7 | Cell Signaling Technology | 12390 |
| Anti-IRF7 | Cell Signaling Technology | 13014 |
| Anti-p-IRAK4 | Cell Signaling Technology | 11927 |

| | | |
|---|---|---|
| Anti-IRAK4 | Cell Signaling Technology | 4363 |
| Anti-MD2 | ProteinTech | 11784-1-AP |
| Alexa Fluor 568-conjugated goat anti-rabbit IgG | Invitrogen | A-1103 |
| Anti-Rabbit HRP-linked IgG | Cell Signaling Technology | 7074 |
| Zombie Aqua Fixable Viability Kit | Biolegend | 423102 |
| Fc Receptor Blocking Ab | ThermoFisher | 14-9161-73 |
| Fluorescent-conjugated antibodies against TLR4 (human) | ThermoFisher | 12-9917-41 |
| Fluorescent-conjugated antibodies against CD14 (human) | Biolegend | 301826 |
| **Oligonucleotides and other sequence-based reagents** | | |
| **PCR Primers** | | |
| Mouse *Hif1α*<br>F: ATT TTT GGA CAC TGG TGG CT<br>R: ATG CAA TGG TGA AAT GCT GA | This study | |
| Mouse *Arg1*<br>F: TTA GAG ATT ATC GGA GCG CCT TTC<br>R: CCG TGG TCT CTC ACG TCA TAC TCT | This study | |
| Mouse *Il1r1*<br>F: TTT ACT CCGAAG AAG CTC ACG TTG<br>R: TGG AGG TCT TGT GTG CCC TTA TGT | This study | |
| Mouse *Cd274*<br>F: GTG AGT GGG AAG AGA AGT GTC ACC<br>R: GGC TGT GAT CTC CAA AAC GTA CAG | This study | |
| Mouse *Marco* | This study | |

| | | |
|---|---|---|
| F: CTT AGC AGC TAT GGA GGT GGC TCT<br>R: ACA CCC GCA TCT TCA TTA TGT ACG | | |
| Mouse *Il-1β*<br>F: GAT CCC AAG CAA TAC CCA AAG AAG<br>R: CTC TGC TTG TGA GGT GCT GAT GTA | This study | |
| Mouse *Tbp*<br>F: GAA TAA GAG AGC CAC GGA CAA GG<br>R: AAG CCC AAC TTC TGC ACA ACT CTA | This study | |
| Human *IFNB*<br>F: GCA CTG GCT GGA ATG AGA CTA TTG<br>R: TCC CCT GGT GAA ATC TTC TTT CTC | This study | |
| Human *GAPDH*<br>F: GTC ATT GAG AGC AAT GCC AG<br>R: GTG TTC CTA CCC CCA ATG TG | This study | |
| **Chemicals, Enzymes and other reagents** | | |
| Human recombinant DKK1 | Biolegend | 778602 |
| Mouse recombinant DKK1 | Biolegend | 759604 |
| MCC950 | ApexBio | B7946 |
| Mouse M-CSF | BioLegend | 576404 |
| LPS | Sigma | L4391-1MG |
| Nigericin | Sigma | SML1779-1ML |
| PMA | Selleckchem | S7791 |
| MSU | Sigma | U2875-5G |
| Nano-$SiO_2$ | Invivogen | tlrl-sio-2 |

| | | |
|---|---|---|
| SN50 (NFκB inhibitor) | MedChemExpress | HY-P0151 |
| NG25 (TAK1 inhibitor) | Selleckchem | S8868 |
| TAK-242 (TLR4 inhibitor) | Selleckchem | S7455 |
| BAY 11-7082 (NFκB inhibitor) | Invivogen | tlrl-b82 |
| GSK8612 (TBK1 inhibitor) | MedChemExpress | HY-111941 |
| Rapamycin (mTOR inhibitor) | Selleckchem | S1039 |
| LY294002 (PI3K inhibitor) | ApexBio | A8250 |
| GSK653094 (SGK1 inhibitor) | Tocris | 3572 |
| TWS119 (GSK3β inhibitor) | ApexBio | B1540 |
| WAY-262611 (DKK1 inhibitor) | MedChemExpress | HY-11035 |
| RPMI | Gibco | 11875-093 |
| DMEM | Gibco | 11960-044 |
| PBS | Gibco | 10010-023 |
| MEM-NEAA | Gibco | 11140-050 |
| Sodium Pyruvate | Gibco | 11360-070 |
| Glutamax | Gibco | 25030-081 |
| Penicillin/Streptomycin | Gibco | 15140-122 |
| Trypsin | Gibco | 2300-054 |
| FBS | GeminiBio | 324000 |
| TRIzol | Invitrogen | 15596026 |
| PVDF membrane | BIO-RAD | 1620177 |
| Paraformaldehyde (PFA) | Thermo Scientific | J19943-K2 |
| 10X Cell lysis buffer | Cell Signaling Technology | 9803S |
| 100X Protease inhibitor | Cell Signaling Technology | 5871S |
| 100X Phosphatase inhibitor | Selleckchem | B-15001 |

| | | |
|---|---|---|
| Phenylmethylsulfonyl Fluoride (PMSF) | Sigma | 10837091001 |
| 5X RIPA Buffer (5% NP-40) | Alfa Aesar | J62524 |
| Protein A Magnetic bead | Cell Signaling Technology | 73778S |
| Disuccinimidyl Suberate (DSS) | ThermoFisher | 21655 |
| Red Blood Cell Lysis Buffer | Biolegend | 420302 |
| PowerUp SYBR Green master mix | ThermoFisher | A25776 |
| SignalFire ECL reagent | Cell Signaling Technology | 6883 |
| CyQUANT™ LDH Cytotoxicity Assay kit | Invitrogen | C20300 |
| Mojosort Mouse Monocyte Isolation Kit | Biolegend | 480153 |
| ELISA MAX™ Deluxe Set Mouse IL-1β | Biolegend | 432604 |
| ELISA MAX™ Deluxe Set Human IL-1β | Biolegend | 437004 |
| GeneJet RNA Purification Kit | Thermo Scientific | K0731 |
| NE-PER Nuclear and Cytoplasmic Extraction Reagents | ThermoFisher | 78833 |
| Pierce™ BCA Protein Assay Kit | ThermoFisher | 23225 |
| 8-well Chamber | ThermoFisher | 154941pk |
| VECTASHIELD Vibrance Antifade Mounting Medium with DAPI | Vector Laboratories | H-1800 |
| High-Capacity RNA-to-cDNA Kit | ThermoFisher | 4387406 |
| Dual-Glo® Luciferase assay System | Promega | E2920 |
| Lipofection™ Transfection Reagent | ThermoFisher | L3000001 |
| **Software** | | |
| FlowJo | FlowJo, LLC | version 10.7, Tree Star |

| | | |
|---|---|---|
| GraphPad Prism | GraphPad Software Inc. | version 9.4.1 |
| **Other** | | |

#### Expanded View Figure 1

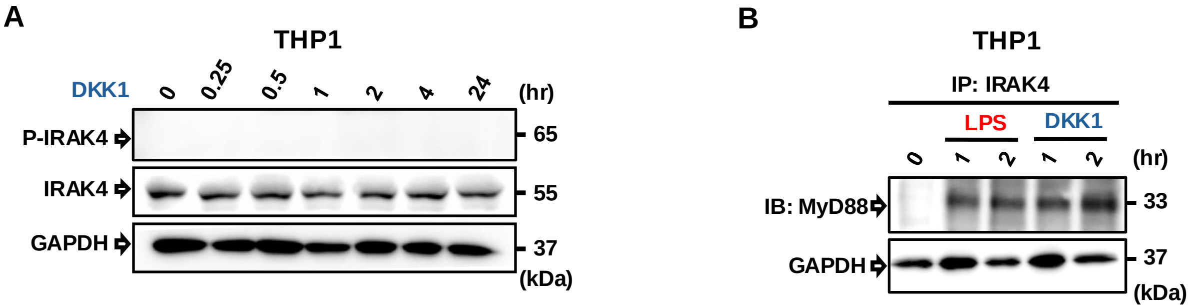

### Expanded View Figure 2

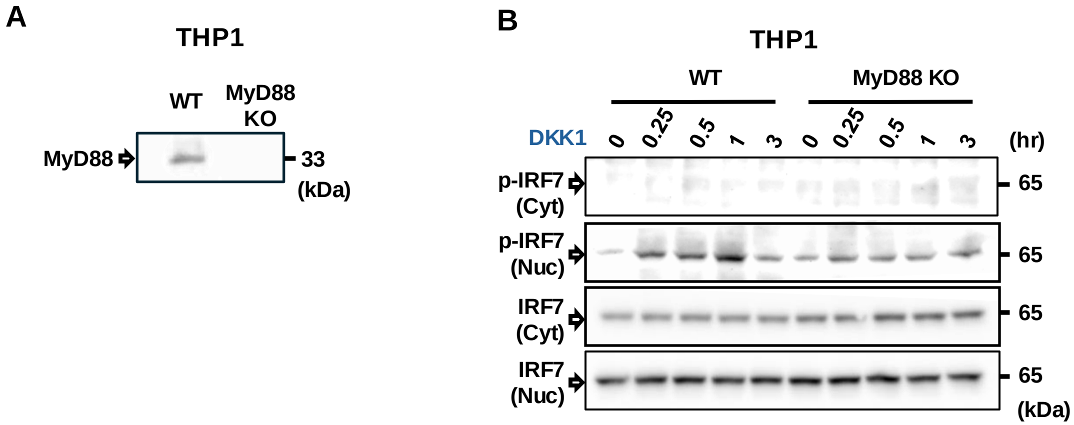

### Expanded View Figure 3

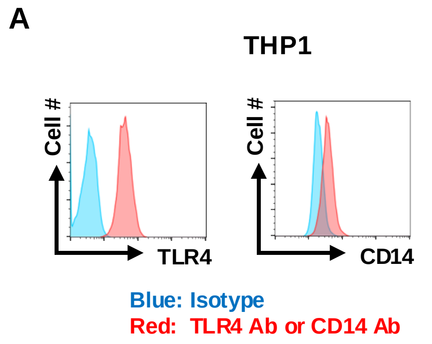

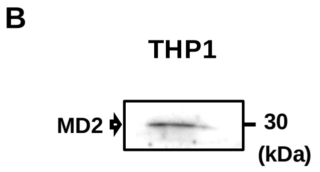

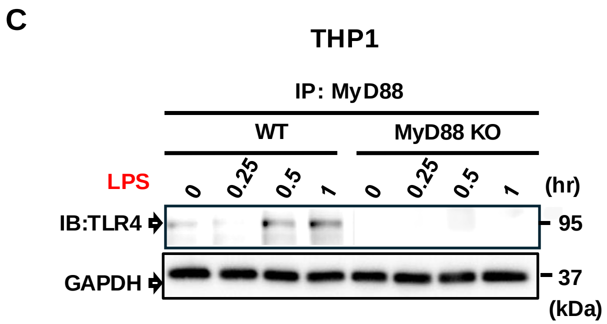

### Expanded View Figure 4

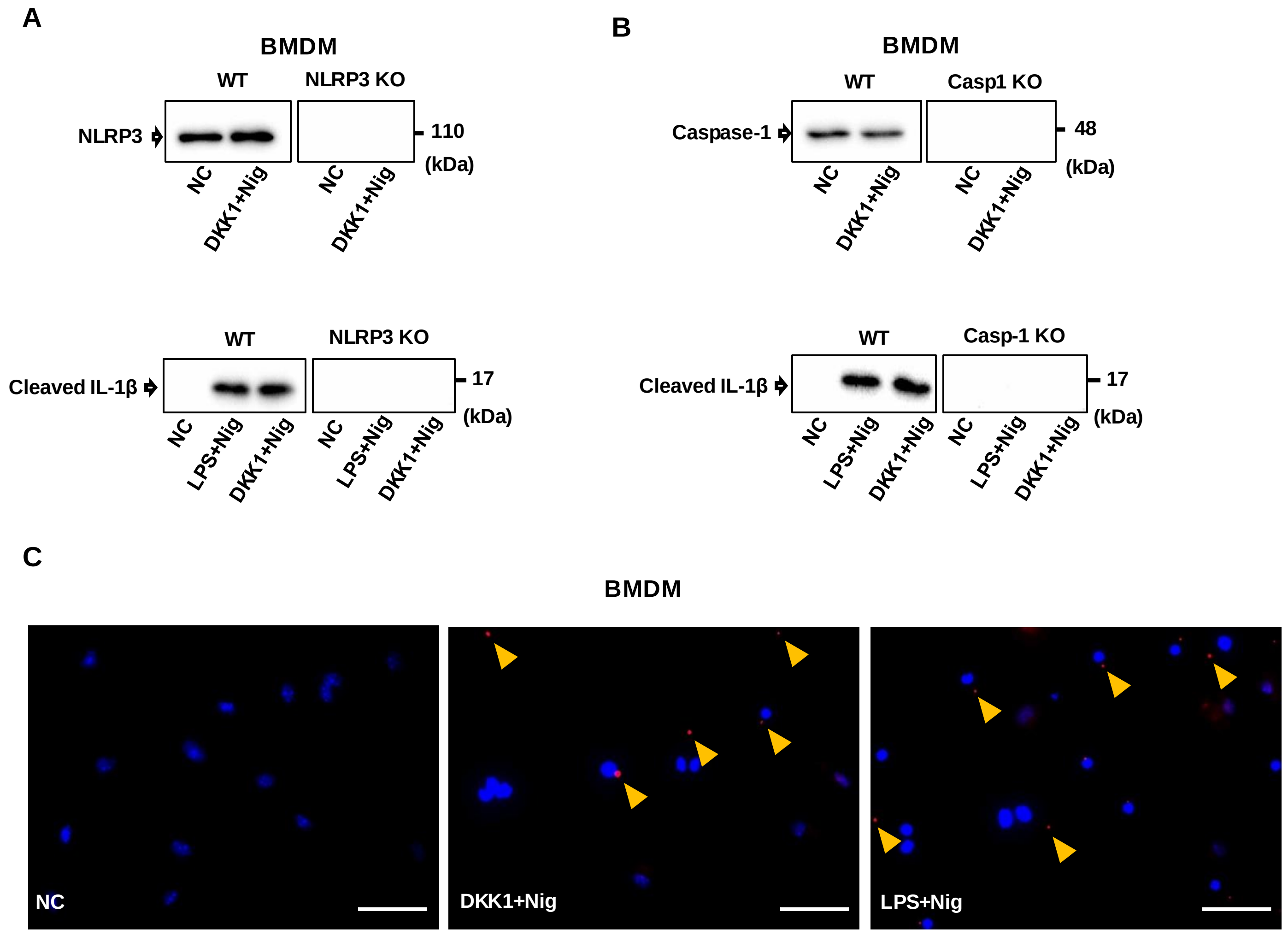

#### Expanded View Figure 5

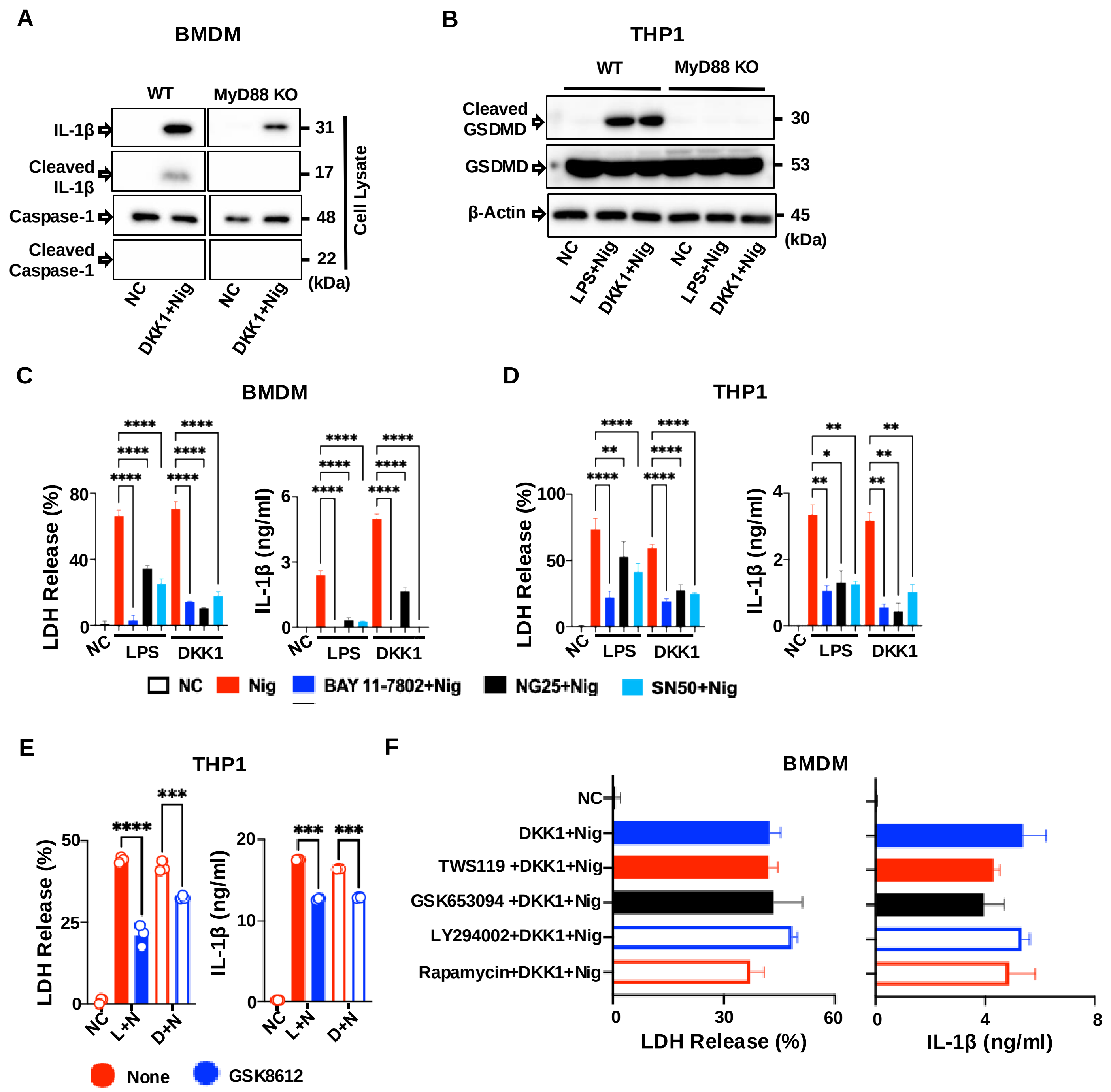
